# Knowledge of theory of evolution and beliefs as determinants of dualism in health science students

**DOI:** 10.1101/2022.02.28.481598

**Authors:** María Alejandra Petino Zappala, Pablo Luis López, Felipe Aguirre, Pablo Richly

**Author notes:** Corresponding author. Postal address: Intendente Güiraldes 2160, Pabellón II. Zipcode: C1428EGA, CABA, Argentina. Declarations of interest: none. Funding support: none of the authors received specific funding for this research.

## Abstract

It has been proposed that dualistic conceptions on the mind and the body can affect the practice of health professionals. Religious beliefs have already been described as affecting mind-body dualistic thinking. Another factor that may play a role is knowledge of evolutionary theory, but this relationship has not been explored. In this work, data on knowledge of evolutionary theory and supernatural and dualistic beliefs of 287 Argentinian students of psychology and medicine majors. were collected through a cross-sectional online survey. Information was analyzed to determine whether an association exists between knowledge of evolutionary theory, supernatural beliefs and dualism. We found significant statistical associations between supernatural beliefs and dualism (positive) and between both and knowledge of evolutionary theory (negative). Also, some heterogeneity was found within dualistic questions between theoretical ones and a hypothetical case. Our results are consistent with the hypothesis that knowledge of evolutionary theory could challenge mind-body dualistic conceptions.

## Introduction

The question of the correspondence between “the mind” and the body has accompanied humanity for centuries and has been a core subject of discussion for religion, philosophy and the sciences. In this work, we will focus on mind-body dualism, a theory or system of thought that interprets reality in terms of two independent principles, in this case mind and matter (i.e. the body / brain). According to a strict dualistic stance, physical and mental properties of individuals are thought to be distinct in their nature and ontology, which holds implications for the understanding of thoughts, consciousness, identity, free will, embodiment, and more practically in the approach to the relationship between mental and physical states, including health problems (see below). While traditionally regarded as a western, classical or modern system of thought, mind-body dualistic conceptions seem to occur transculturally in our species, as some degree of dualistic thinking has been found in people of different societies, and in pre-modern societies (Chudek et al., 2018; Eccles, 1989; Slingerland & Chudek, 2011).

Conversely, different hypotheses link the brain and/or the body to phenomena of the mind, which range from reductionist explanations of the mind as entirely a manifestation of neuron activities, to multilevel materialistic explanations, to others defining mental and physical phenomena as metaphysically different but still connected (Stoljar, 2022). While the issue is far from resolved, a vast amount of empirical findings from research in neuroscience shows mensurable correlates to thought processes, challenging strict “substance” dualism. Nevertheless, it is not clear that this evidence is sufficient to fully understand thought, consciousness or psychopathology (Moreira-Almeida, Araujo & Cloninger, 2018; Motuca, 2016). In fact, regardless of the advancement of neurosciences, the dualist stance is frequently observed in the general population, including scientists and health professionals (Strejilevich et al., 2014). Moreira-Almeida, Araujo & Cloninger (2018) contend that while literature on psychiatry and the neurosciences mostly adheres to materialism, the bias towards simplistic and reductionist explanations makes some questions on consciousness intractable. In this sense, the prevalence of dualistic beliefs in health professionals may reflect a dissatisfaction with the limits of reductionist approaches to consciousness and related phenomena (Moreira-Almeida & Araujo, 2015). On the other hand, Mehta (2011) argues that the persistence of a dualistic mindset in mental health professionals can be explained because mind and body dualism is a convenient philosophy that uses the “divide and conquer” strategy to cope with prevalent religious thinking, and would be, therefore, more convenient to deal with the complexity of human nature.

In the case of those who are dedicated to mental health, determining the extent and the effects of this belief is critical, since the conceptions about the mind can affect professional practice (Fahrenberg & Cheetham, 2000). Believing that consciousness and the body do not correspond in any way to each other can become, for example, an obstacle to the diagnosis of psychiatric symptoms secondary to physical disorders, can result in the underestimation of physical interventions on mental health, or may lead to consider the patients as responsible or blameworthy for their ailments (Glannon, 2020; Miresco & Kirmayer, 2006). Moreover, a strict dualistic stance may predispose these professionals to ignore or downplay the importance of material conditions of existence, the effects that cultural practices can have in the shaping of the body and subjectivity, among other problems that have been raised by critical scholars (Karmakar, 2021; Lee, 2014). That is why it is vital to be able to improve the understanding of the factors affecting or determining this philosophical stance in order to design strategies aimed at revising dualistic conceptions during the formative period of future health practitioners.

Some factors have already been described as important in determining the likelihood that a person adheres to this system of thought. Religiousness has been proposed as such; while the idea of afterlife and an immortal soul, prevalent in theists, does not necessarily imply a mindbody dualistic stance (Cottingham, 1992), a stronger religiousness is frequently associated with this position, even among health professionals (Demertzi et al., 2009). On the other hand, it has been argued that exposure to neuroscientific accounts of consciousness phenomena could decrease mind-body dualistic beliefs (Harrington, 2013), but conversely, an emphasis on the limitations of these explanations may predispose people to take a dualistic stance or, at least, to question a reductionist mechanistic approach (Moreira-Almeida & Araujo, 2015; Preston et al., 2013). It should be noted that the last two studies constitute experiments in which the effect of an exposure on dualistic thinking was assessed shortly after the event and may not reflect long term effects. Moreover, exposure may have heterogeneous effects depending on previous beliefs (Harrington, 2013).

Historically, it has also been proposed that the emergence of the field of evolutionary biology has undermined the basis of mind-body dualistic thought in the natural sciences (Costall, 2004; Searle, 2013), although it is not clear whether it is because of the idea of the mind as a product of natural selection, which can be debated (Lycan, 2009), by undermining the notion of human consciousness as unique or essentially different from other species’ (see below) or because of the materialistic assumptions guiding evolutionary biology. The relationship between evolutionary theory and religiousness has been thoroughly studied: a certain correlation between knowledge about the theory of evolution and religious beliefs in students entering university has already been described (Moore et al., 2011) as well as a small effect of religiousness in the acceptance of evolutionary theory for students of different majors (Gefaell et al., 2020), but it is not known for sure if this variation in evolutionary knowledge has any effect on the likelihood that a person holds a mind-body dualistic perspective independently of their religiousness or, at least, if both are correlated. It has been argued that evolutionary theory encompasses a materialist theory of the mind, in the sense that it is seen as an emergent of the physical world and subject to similar laws or principles as other bodily functions or properties, including of course natural selection (Callinicos, 1996), and in this regard evolutionary theory would be incompatible with a strong mind-body dualistic stance. Moreover, by considering humanity as part of a continuum of life forms, Darwinian theory presents human thought processes as a not essentially distinct from that of non-human animals. This conception also arguably runs counter to strict Cartesian mindbody dualism (Allen & Trestman, 2020). In this sense, presenting students with the concepts and problems concerning evolutionary biology could, in principle, serve to call into question a strict dualistic perspective. Indeed, in the case of a prevalent philosophical stance such as mind-body dualism, making assumptions explicit and offering challenging views can be critical to “unblind” future health professionals to possible biases that may, unbeknownst to them, profoundly affect their conceptions and practices (Mehta, 2011). In this sense, it can be argued that knowledge about the theory of evolution could serve to challenge mind-body dualistic conceptions in future health professionals in a crucial period of their training. It has been indeed suggested that, when presented with theories contradicting intuitive concepts, students must “un-learn” those previous ideas in order to grasp the new information (Shtulman & Harrington, 2016), or at least can learn to “circumvent” those intuitions through metacognitive reflection (González Galli, 2019b), and to this end, presenting them with concepts and problems pertaining to evolutionary theory could make them question, or at least recognize, dualistic assumptions.

To our knowledge, although it would be reasonable to hypothesize that understanding of evolutionary theory could undermine strict mind-body dualistic thinking, no studies have been carried out to evaluate if such association exists. In the present work we tried to answer this question by investigating the relationship between dualistic conceptions, religious or supernatural beliefs and knowledge of the theory of evolution (general and applied to human evolution) in Argentinian students of psychology and medicine majors.

## Materials and Methods

In order to gather the information for this study, we conducted a cross-sectional online survey using an incidental sampling technique.

### Study and questionnaire design

Questions about supernatural beliefs (in God, the soul, energies, afterlife), specific religious affiliation and practices were designed following previous surveys performed in Argentina (CEIL - CONICET, 2019).

Three questions to rate dualistic thinking were included, two of them theoretical, to account for the agreement of the interviewees with dualistic statements, and a hypothetical case scenario to rate their responses to a practical problem (Author, 2022).

Knowledge of Evolution Exam (KEE) questions were translated to Spanish from the original questionnaire (Cotner et al., 2010) and an extra question was added regarding evolutionary mechanisms besides natural selection. As a strong dualistic stance may imply the belief that humans and non-human animals are qualitatively different in their properties (Allen & Trestman, 2020), additional questions were designed for a Knowledge of Human Evolution Exam (KEEH) to account for knowledge of evolutionary theory pertinent to the human species.

All questions were previously validated in a preliminary study using an independent sample not included in the present analysis. The complete questionnaire is provided in Additional Files 1 and 2.

### Data collection and ratings

Online surveys were performed by means of a Google Forms file sent to students of Psychology and Medicine from different Argentinian Universities, either through public social media posts or by means of direct contact (email) with university teachers. The data were collected during November 2020 (Figure 1). The questionnaire included personal data (age, gender, major and university) in addition to Supernatural beliefs, Dualism, KEE and KEEH questions. A target sample size of at least 250 people was established, based on the preliminary studies, to reach adequate statistical power.

**Figure 1.**
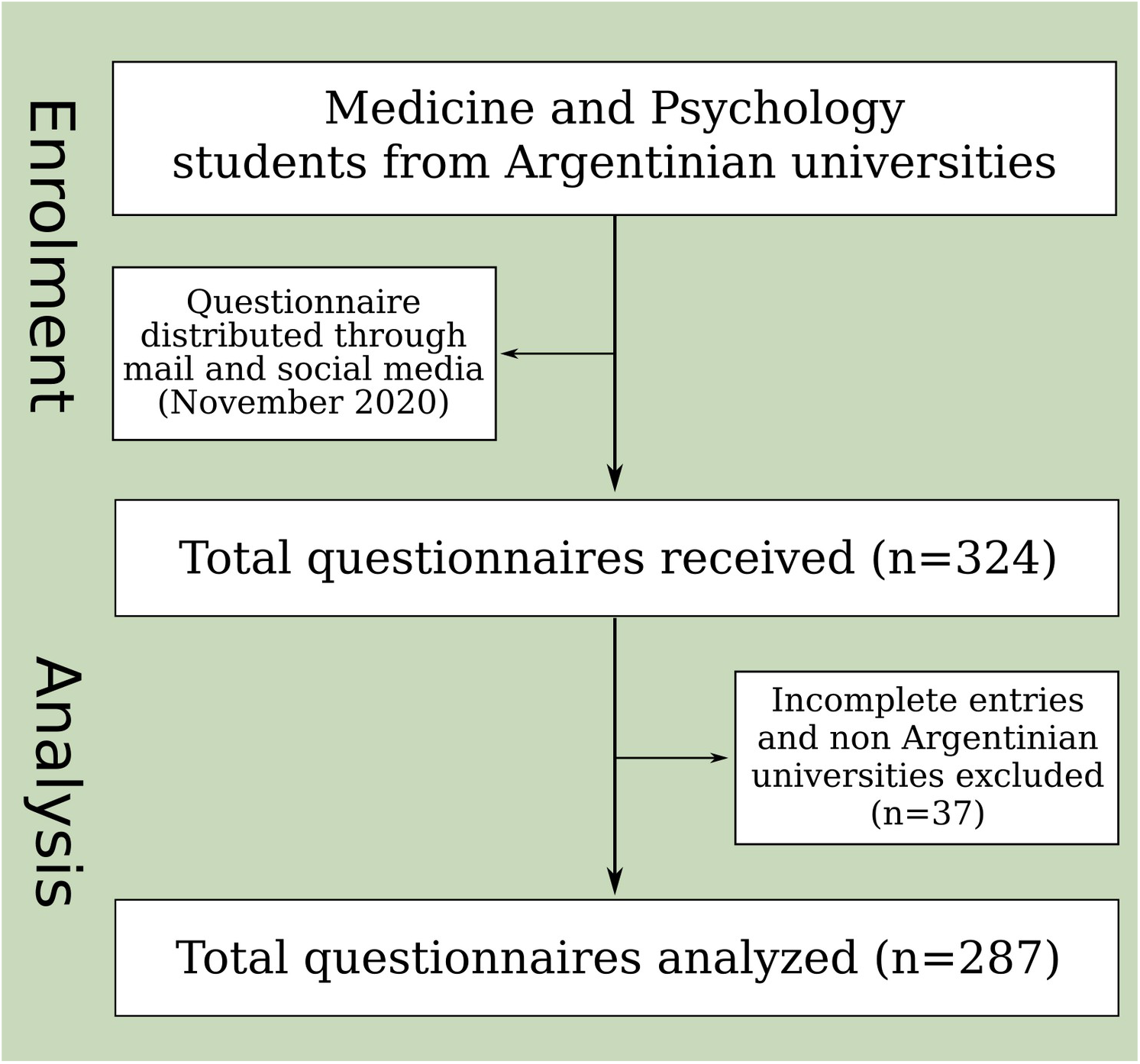
Flowchart corresponding to the survey questionnaire process.

Correct KEE and KEEH answers were rated with a score of 1, with 0 points added for incorrect answers. Only KEE question #11 had three possible outcomes, depending on the interviewee’s selection of all (2 points) or only one of the correct answers (1 point).

Questions on religion and supernatural beliefs were rated from 0 (does not believe in God / does not have any relationship to God) to 2 points (does believe in God / takes part in religious activities or institutions), except the binary question about the religion (where “no religion” amounted to 0 points and the rest of the answers were rated 1).

A similar criterion was followed for questions on dualism, from 0 points (disagree / Juan’s family) to 2 (agree / María’s family).

Overall ratings for KEE, KEEH, Supernatural beliefs and Dualism were obtained by summing all scores for each set of questions.

### Statistical analyses

All statistical analyses were performed in Rstudio (Allaire, 2012) using R version 3.6.3 (R core team, 2020).

First, we wanted to explore whether there were any differences between majors for the answers to the survey, as homogeneity would be an important assumption when pooling the answers for Psychology and Med students. Mann Whitney Wilcoxon Tests were performed for each of the primary outcome scores to evaluate whether students for both majors differed in their religious or dualistic beliefs or in the performance for both Evolutionary Biology knowledge tests. Chi squared tests were also performed for the individual questions to test if the proportions obtained for each possible answer depended on the major.

To inspect the statistical relationship between knowledge of evolutionary theory, supernatural or religious beliefs and dualistic thinking, bivariate Spearman correlations were calculated for KEE, KEEH, Supernatural beliefs and Dualism scores using the rcorr function from the Hmisc R package (Harrell, 2019).

To further study the relationship between variables, in this case in the multivariate space, a Principal Component Analysis (PCA) was performed with the standardized scores for KEE, KEEH, Supernatural beliefs and Dualism using R prcomp function. The data on interviewee’s age and major were later superimposed on biplots but were not included on the PCA analysis as active variables.

The PCA scores were used to test whether individuals studying different subjects occupied distinct sectors of the multivariate space. The scores for individuals in all PCA components were subjected to a permutational multivariate analysis of variance (PERMANOVA, (Anderson, 2001)) using the adonis function of the vegan R package (Oksanen et al., 2020), having previously checked the assumption of multivariate homogeneity of variances (using functions betadisper and permutest from the same package).

Finally, in order to model the probability of answering correctly or incorrectly based on major, supernatural beliefs and gender, logistic regressions were performed for the score on each of the KEE and KEEH questions using the glm function for the binomial family from the R package nlme (Pinheiro et al., 2021). To test the relationship between major, religion and gender and dualistic thinking, logistic regressions were also run for questions on Dualism. In these cases, the multinom function for the nnet package (Ripley & Venables, 2021) was used, as these questions had multiple possible answers.

### Transparency, openness and ethical considerations

We provide information on the determination of sample size, exclusion of cases and data analyses (see above). The complete set of data is provided as supplementary information and analysis code for this study is available by emailing the corresponding author. Information on software and packages used is provided above. This study’s design and its analysis were not preregistered.

This research was conducted following the pertaining ethical guidelines. All survey participants were informed regarding its objectives and declared they agreed to participate. No data were collected that could reveal the interviewees’ identities, therefore the study was not submitted for approval to institutional review boards.

This manuscript follows the Recommendations for the Conduct, Reporting, Editing and Publication of Scholarly Work in Medical Journals, including sections II A, II B, I E, III B. Section III K is not applicable.

## Results

A total of 287 subjects meeting the inclusion criteria answered the survey completely (of which 133 were Med students and 154 were Psychology students; Additional File 3). The study sample included students from 18 different Argentinian universities from various provinces; public and private institutions were represented in the sample. 68% of the respondents were female, 31% male and 1% other, and the mean age was (24.16 ± 5.27) years (mean ± SD).

There were no significant differences between Medicine and Psychology students regarding the primary outcomes (Tables 1 and Additional File S4). Of all the items in the questionnaire, only the answers to 2 questions from the KEE differed statistically for both majors in a Chi squared test, and for only one of them (KEE #6) the test was strongly significant, with p <0.01 (Additional File 4).

**Table 1.** Primary outcomes for the measured scores (mean score and n for each major)

|  | **Supernatural beliefs** | **Dualism** | **KEE** | **KEEH** | **n** |
| --- | --- | --- | --- | --- | --- |
| Medicine | 4.33 | 1.44 | 7.48 | 1.65 | 133 |
| Psychology | 4.18 | 1.45 | 7.25 | 1.52 | 154 |

Beliefs in the existence of a supernatural energy and on a soul were more prevalent among the interviewed students than beliefs in God or in afterlife (Figure 2), but all items were less frequently reported in our sample than in the general population from Argentina (CEIL - CONICET, 2019). We found that the proportion of people who believed in the existence of the soul was higher than those who believed in God or an afterlife, but similar percentages were observed for the questions about the belief in afterlife and the independence of the soul and the body, indicating that some may believe in a soul rooted in a material substrate (i.e. not substance dualists). However, the soul-body dualistic conception was still the most prevalent dualist answer, and the only one where disagreement was the minority option (Figure 2). On the other hand, when the question was framed in terms of the independence between mind and brain, more than 80% of interviewees disagreed.

**Figure 2.**
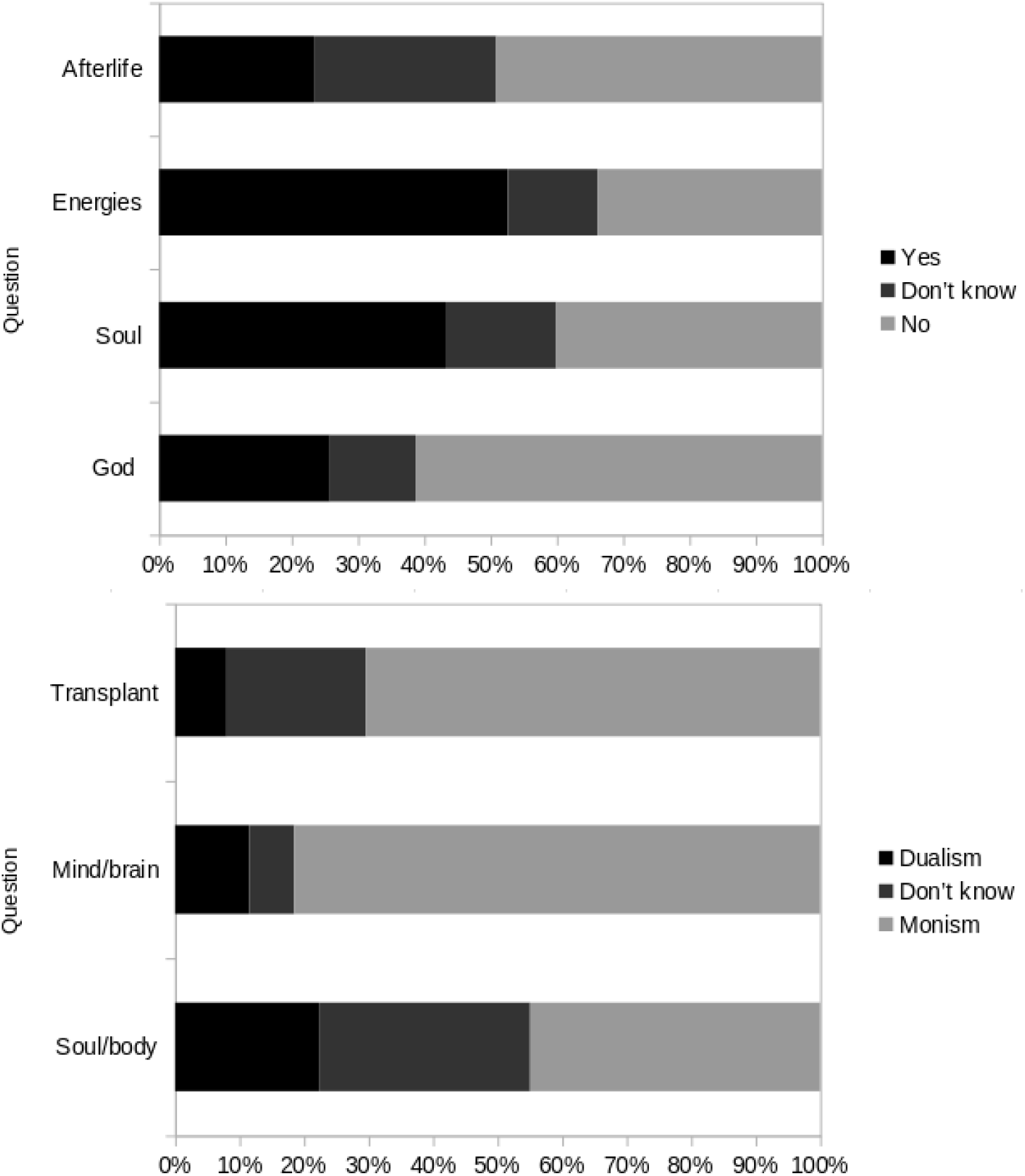
Responses about supernatural beliefs and dualism. Percentages for each answer are depicted.

Regarding the correlations between the primary outcomes (Table 2) we found a positive linear relationship for Dualism and Supernatural belief scores, as described in previous reports (Demertzi et al., 2009), and a negative correlation of both with KEE score. However, only Supernatural beliefs showed a significant negative association with KEEH score. Unsurprisingly, both Knowledge of Evolutionary Theory scores presented a positive correlation with each other, although it was weaker than expected.

**Table 2.**
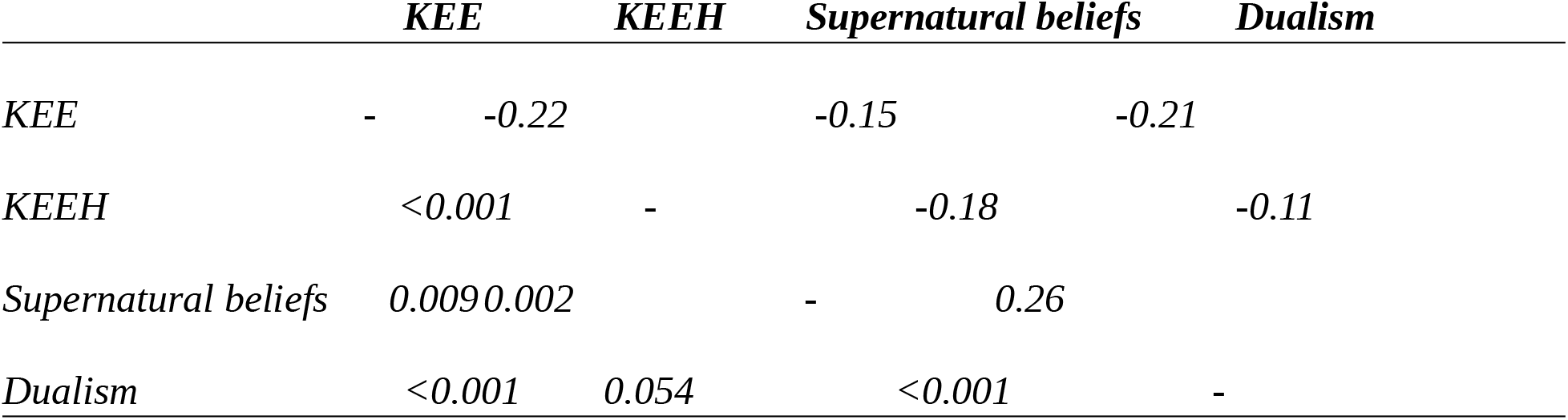
Spearman correlations for the primary outcomes scores (above the diagonal); p-values below the diagonal.

To better understand the multivariate relationships of the measured scores we performed a Principal Component Analysis, in which information about the initial variables (individual scores obtained) is ordered and summarized in new components, and a space is created with these components in which individuals are sorted in relation to their answers (with individuals answering similarly located near each other). The first two components explained 61% of the total variance (Additional File 5). Component 1 correlates positively with Knowledge of Evolution scores and negatively with Beliefs and Dualism scores. Component 2 shows that, regardless of the main negative tendency between these groups of scores, individuals with high KEE scores and high degree of Dualism and Beliefs, and vice-versa, can be found in the sample. In general terms the relationships between KEE, KEEH, Supernatural beliefs and Dualism reflect what was found in previous analyses and no apparent pattern was found regarding age or major of the interviewees (Figure 3).

**Figure 3.**
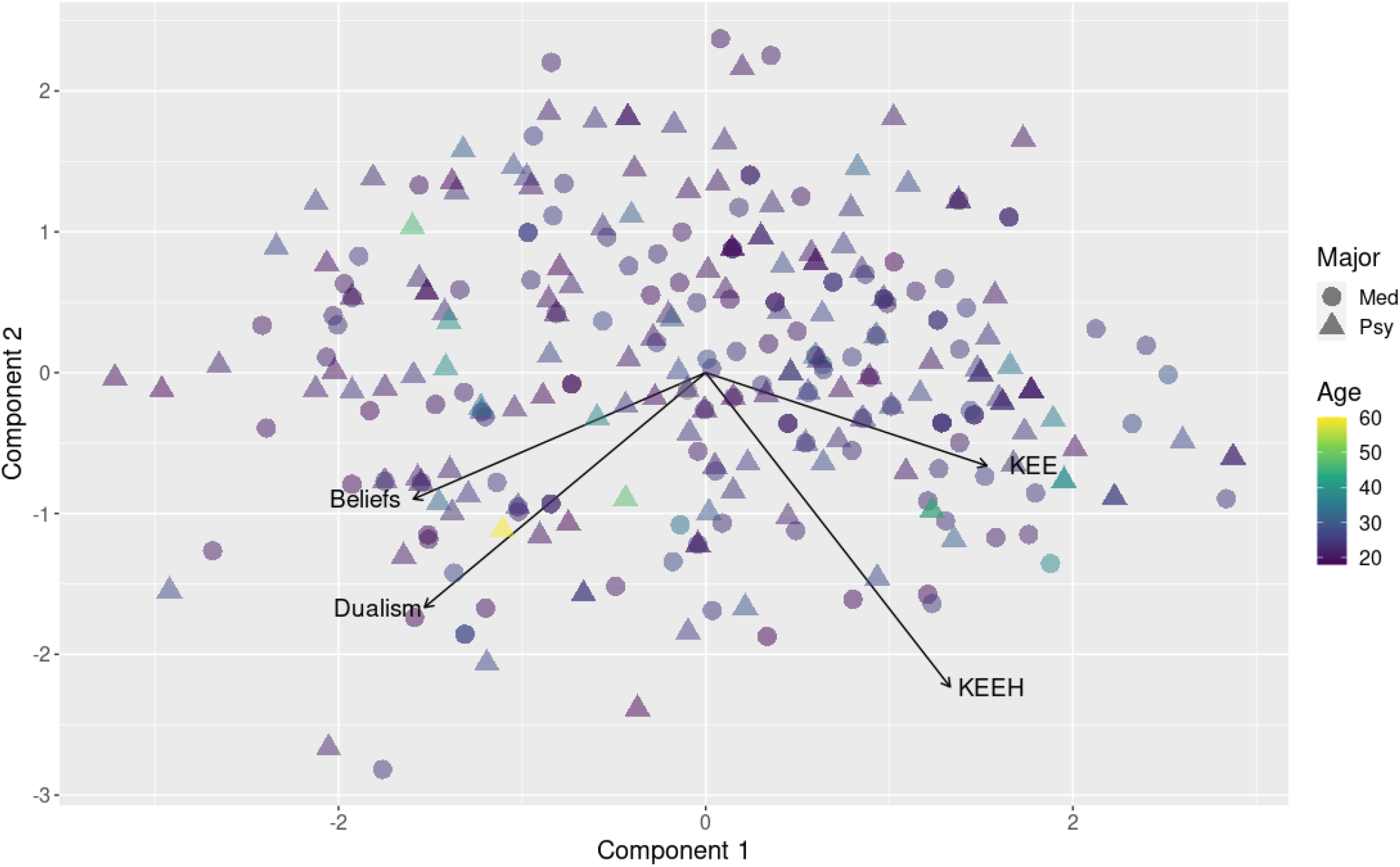
Results for the first two components of the PCA for Dualism questions. The shape of each data point represents the major and the color scale indicates participant’s age

However, to further confirm our previous findings that students for both majors do not differ significantly in their responses, but in this case in the multivariate space, the scores for all individuals in all PCA components were used for a permutational analysis of variance (PERMANOVA), after confirming no significant departures from homogeneity of multivariate dispersion in this space. Multivariate scores for both majors did not differ (p>0.05), meaning the students for both Medicine and Psychology do not occupy different sectors in the space (i.e. the groups do not differ significantly in their scores).

It is worth noting that as we assessed the answers to Dualism questions separately with another PCA consisting only of these scores, we found some degree of heterogeneity, given that the responses to theoretical questions apparently behaved independent from the hypothetical case scenario. In this case it can be seen that the first component correlates with mind/brain and spirit/body dualistic scores, while the hypothetical case scenario scores was not an important variable defining this component. Conversely, the second component correlates strongly with the scores for this case scenario (Figure 4). Extracting this item from the analysis, however, did not significantly modify the main correlations, so this finding would not be relevant for the present analysis, although we find it interesting and worthy of further study, as we believe this heterogeneity may be mediated by different types of dualism in the sample.

**Figure 4.**
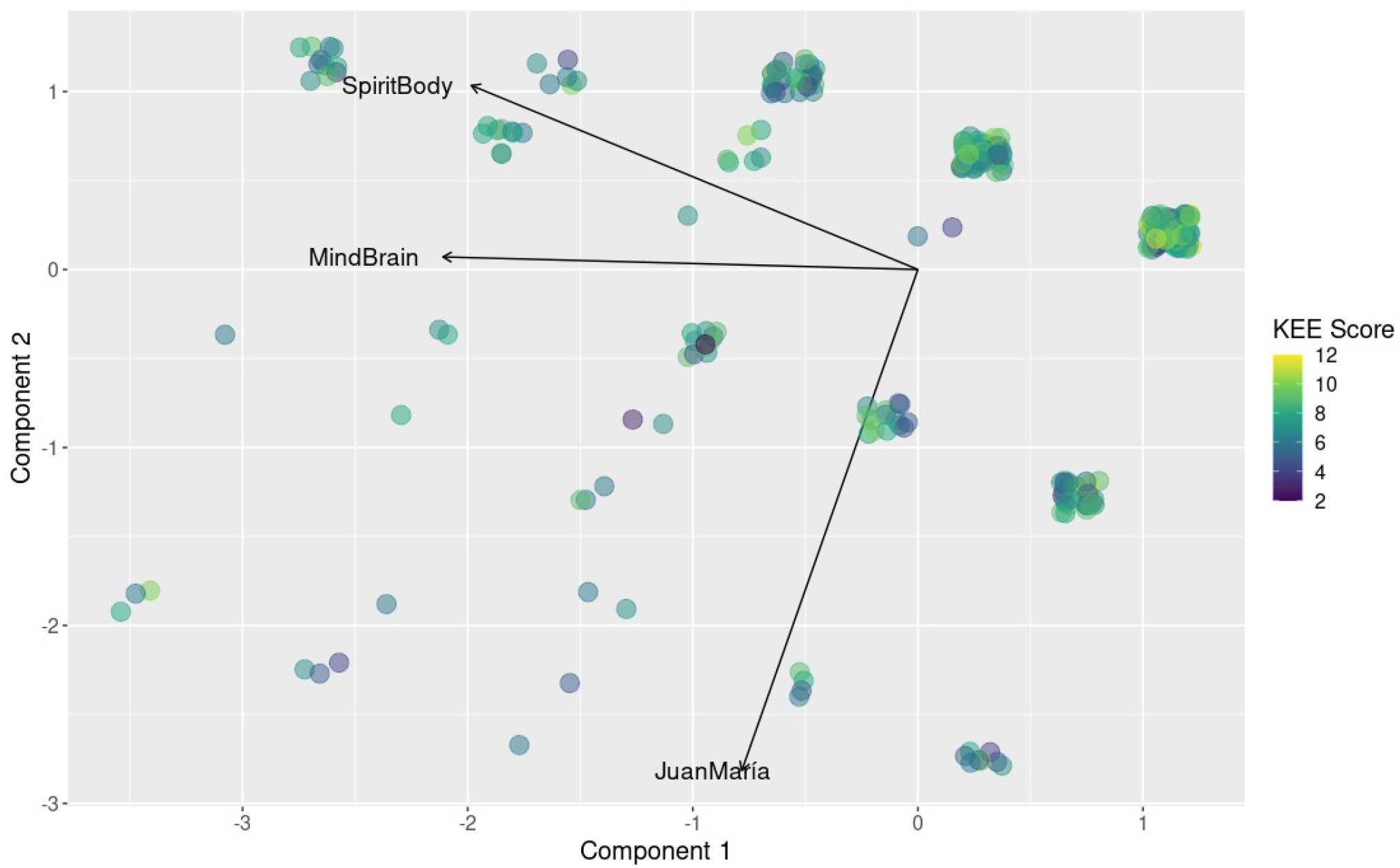
Results for the first two components of the PCA for Dualism questions. The color scale indicates participant’s performance on the KEE questionnaire.

Finally, logistic regressions were performed in order to model the probability of answering correctly or incorrectly to KEE and KEEH questions, based on major, religion and gender (Additional File 6). In most cases, only religion showed a significant effect on the likelihood of a correct answer for KEE questions (where nonreligious participants performed better than religious people). No effects of either religion, gender or major were found for KEEH answers.

On the other hand, on the Dualism question regarding the relationship between body and spirit, nonreligious people were more likely to answer agnostically (“I don’t know”) or to deny the existence of such relationship. They also tended to answer against the dualist option in the hypothetical case scenario. In the question regarding mind and brain, men were less likely to answer “I don’t know” than women.

## Discussion

Mind-body dualism is a widely shared system of thought that, as we mentioned, might profoundly impact the understanding not only of human nature but also of our interactions with other species and the environment (Le Grange, 2004). While it may have been useful to deal with some degree of complexity in understanding natural phenomena, including health, it can be argued that today a strong substance dualism represents an obstacle for those working in these areas (Mehta, 2011). This philosophy may entail the neglect of material conditions of existence and of the body itself as affecting the psychological state and well-being, and therefore preclude a comprehensive understanding of health issues. Consequently, some authors argue that dualism can exert negative influences on the practice of health professionals, especially (but not limited to) those observing mental health. As has been established before, this stance has been criticized and questioned because it may obscure how gender, race and class issues shape our body and health (Karmakar, 2021; Lee, 2014). In fact, the relevance of strong dualism as a problematic philosophy in future health professionals appears more important when considering the evidence that the more a behavioral problem is seen as originating in psychological (i.e. “not physical”) processes, the more a patient tends to be viewed as blameworthy for their symptoms; conversely, the more behaviors are attributed to physical or biological causes, the less likely patients are viewed as responsible (Miresco & Kirmayer, 2006). Therefore, according to these authors, a dualistic approach might lead to a voluntaristic view regarding mental and other health issues that would make patients accountable of their own suffering, while simultaneously neglecting the critical aspect of the social and material conditions affecting health. Moreover, this conception might be an obstacle not only to practice, but also to health research. Mehta (2011) argues that Cartesianism is usually associated to methodological implications such as reductionism and a rigid positivism, ignoring subjectivity and leading to a crisis of the field. Bunge (2008) also contended that a dualist approach defined a separation of the biological and the psychological that precluded the understanding of mental evolution. On the other hand, criticism to neuroscientific reductionism and the inability of molecular explanations to account for different conscious phenomena has been raised as a reason for the belief on a mind or soul as separate from the body (Preston et al., 2013). In addition, it has been argued that reductive physicalism can also be a source of stigmatization (Moreira-Almeida, Araujo & Cloninger, 2018), that exclusive reliance on molecular explanations of conscious phenomena may obscure social causes of psychic suffering to favor economic interests of the pharmaceutical industry (Tabb, 2021). For these reasons, and assuming that adherence to any of these stances may affect professional practice, understanding the factors affecting or determining the notions about mind-body dualism in future health workers, as well as identifying devices that foster a conscious reexamining, is of great interest.

The present study has the peculiarity of analyzing the association of both supernatural beliefs and knowledge of the theory of evolution with mind-body dualism in Argentinian undergraduate students from health-related majors, something that, to our knowledge, has not been assessed before. Our most relevant findings show that supernatural beliefs have a positive correlation with dualism, while knowledge of the theory of evolution shows a negative relationship. However, although they are significant, the magnitude of these correlations is weak, probably owing to the heterogeneity present in the sample and the fact that different types of dualism may be confused in the answers. In favor of the latter interpretation, the different proportions of interviewees that believe in the existence of a soul, in its independence of the body and in the dualism of the mind and the brain signal the multiple dimensions of dualistic thinking that may be left unresolved in this questionnaire.

Although no significant differences were found between majors for most of the answers, the composition of Psychology and Medicine programs is, as expected, different. Particularly, Argentinian universities show a strong bias towards psychoanalytic content and no mandatory training in Neurology for Psychology degrees (CONEAU [Comisión Nacional de Evaluación y Acreditación Universitaria], 2009; Fierro & Menéndez, 2020; Mustaca & Franco, 2018). However, contents on evolutionary theory are marginal in the programs for both majors (CONEAU [Comisión Nacional de Evaluación y Acreditación Universitaria], 1999, 2009).

We are aware of the limitations of our study. First of all, no information was collected regarding the degree of career advance for each participant. Biases inherent to voluntary surveys could exist, although the most obvious factors (gender, age) are already biased for students from both majors with respect to the general population (Ministerio de Educación de la Nación Argentina, 2019), showing a composition similar to that of our sample. We also found a bias towards less dualistic conceptions and a lower prevalence of supernatural beliefs that in the general population, a finding that could actually accurately reflect the situation of undergraduate students (Guest & Sharma, 2011; Hayes, 2000), and could generally be regarded as a common occurrence for health-related majors (Gross & Simmons, 2009), but that could cause correlation analyses related to dualism to be underpowered. On the other hand, our sample included students from both majors from different universities in the country, covering several provinces. It encompassed a wide range of scores for all primary outcomes, and both subsamples, obtained by two different methodologies (formal email recruitment by university teachers and through public social media posts) showed similar results. These facts, and the information stated in the previous paragraph, led us to believe that the sample here analyzed adequately represents the population of Argentinian undergraduate students for these health-related majors.

Another downside of this analysis is the exclusively quantitative nature of the data; in this sense, this work could provide for a starting point to design future surveys which could include qualitative questions, as mixed approaches may be better suited to capture the heterogeneity of the interviewees’ knowledge and beliefs. As previously mentioned, a more detailed questionnaire could also help differentiate between different types of dualism which are not distinguishable in the present survey (Moreira-Almeida, Araujo & Cloninger, 2018).

Finally, the design of the study does not allow us to shed light on the causal nature of the associations found, although the results support the hypothesis that both religious beliefs and knowledge of evolutionary theory could affect mind-body dualistic thinking. Future studies could be carried out to verify this relationship with repeated surveys before and after debates on evolutionary biology; this would not only help determine causality but would also be useful for students to acknowledge the changes of their own beliefs, which are frequently overlooked (Preston et al., 2013). Moreover, it would be desirable to reassess in future works the relationship between dualistic and physicalist stances and professional practice, in which we based our interest for this study but that was not evaluated here.

Other studies have found religiousness and supernatural beliefs as positively associated to dualism (Dawkins, 2008; Demertzi et al., 2009; Tayeb et al., 2017), although the causality of the relationship is not clear. It is important to note that freedom of thought, conscience and religion constitutes an inalienable human right (United Nations, 1949), and of course no intervention should be considered in such direction. However, given the similarly significant negative correlation we found between knowledge of evolutionary theory and dualistic beliefs, it may be considered a potentially important factor. Although a correlation does not imply causation, the core assumptions of evolutionary theory and the biological sciences in general, at least in their present form, imply a continuum between the human species and the rest of the living beings, with a strong focus on the material conditions of existence and with consciousness considered as an emergent property of bodily substrates. As such, it has been argued that there is a certain degree of incompatibility between mind-body dualism and knowledge of evolutionary theory (González Galli, 2019a). Moreover, contemporary evolutionary biology tends to offer a variety of explanations that avoid reductionsm, in which the obvious limitations of assigning complex processes exclusively to molecular phenomena can be overcome.

Although we did find results consistent with a negative relationship between knowledge of evolutionary theory and mind-body dualism, our work does not clarify how it arises; it is likely that both factors affect each other. Strong mind-body dualism, when not challenged, may pose an obstacle to the understanding or acceptance of evolutionary theory, but it is also possible that the reflection on the nature of evolutionary change provides the students with a challenge to these dualistic assumptions. Indeed, learning often entails a restructuring of knowledge and concepts, particularly when new information contradicts previously held notions (Blancke et al., 2012). A conscious examination of the assumptions behind different epistemological obstacles to conceptual change is a requisite to achieve it (Shtulman & McCallum, 2014), and this reflection may be triggered by exposing the students to data that contradict their cognitive biases (González Galli, 2019b). While in fact the incompatibility does not preclude the coexistence of evolutionary biology knowledge and dualistic and/or supernatural beliefs in separate domains of thought, providing explanations to different sets of phenomena (Short & Hawley, 2015; Shtulman & Lombrozo, 2016), reflection on evolutionary theory could imply the questioning of strong dualistic assumptions affecting future medical practice of undergraduate students and, at least, promote a conscious examination of such assumptions. Indeed, as Mehta (2011) argued, it is crucial that health professionals are aware of the philosophical framework within which they operate and how it affects their understanding of health issues and their practices; the high prevalence of mind-body dualism and the fact that it is not usually made explicit imply that these cognitive biases are never challenged nor made conscious, and the same would be important for reductionist explanations, arguably of common occurrence in some fields of the health sciences.

It has been argued that intuitive concepts, even when presumably abandoned, can still resurge and cause cognitive conflicts, something that happens even to expert scientists (Shtulman & Harrington, 2016), but this occurs in certain conditions such as during quick reactions. This information suggests that intuitive patterns may not be completely eliminated, but it has been proposed that they can be circumvented through cognitive reflection (i.e. deliberate reasoning) (González Galli, 2019b; Shtulman & McCallum, 2014). The formative period may, indeed, be critical to the reflection on such prevalent notions as mind-body dualism (Mehta, 2011). In this sense, Roselli, Castellaro & Peralta (2022) argue in favor of the promotion of “socio-cognitive conflicts”, i.e. fostering debates in which their multiple perspectives might make previous beliefs enter in conflict, allowing them to transcend their own thinking schemes. Debates on evolutionary theory, on its meaning for the understanding of human as embedded in nature, and consciousness or thought processes as affected by evolutionary processes, may fulfill this role in the classroom. Therefore, we consider our findings as a rationale to propose the inclusion of evolutionary biology in the study plan of those fields of study related to the care of patients suffering from both physical and mental health conditions, although the efficacy of this intervention in professional practice should be confirmed in prospective trials.

## Acknowledgments

We thank all survey participants and all people who shared the request for volunteers.

## Data availability

All relevant data used are available as Supplementary Files.

