## Additional File 1 for "Knowledge of theory of evolution and beliefs as determinants of dualism in health science students"

Additional File 2. Additional KEE and KEEH questions, religion and dualism questions

11 [KEEH1]. According to evolutionary theory, the present form and features of human beings is best explained as:

A. a result of chance only

B. a mystery

C. a creation by a superior intelligence

D. a miracle

E. none of the above is correct

12 [KEEH2]. According to evolutionary theory, modern human beings:

A. are the most evolved species

B. are not subject to natural selection anymore

C. are superiors to their ancestors

D. more than one of the above is correct

E. none of the above is correct

13 [KEEH3]. According to evolutionary theory, chimpanzees:

A. are still evolving to become more like human beings

B. are humans’ ancestors

C. share a common ancestor with humans

D. more than one of the above is correct

E. none of the above is correct

14 [KEE11]. Biological evolution:

A. can occur as a result of chance

B. can occur without phenotypic changes

C. can result from migratory movements

D. all of the above are correct

E. none of the above is correct

**Religion:**

1. Do you believe in God?

A. Yes

B. No

C. I don’t know

2. Do you believe in the soul?

A. Yes

B. No

C. I don’t know

3. Do you believe in energies?

A. Yes

B. No

C. I don’t know

4. Do you believe in the afterlife?

A. Yes

B. No

C. I don’t know

5. Which is your religion?

A. Christian

B. Other

C. no religion

6. How do you relate to God, mainly?

A. by myself

B. church / temple / community

C. I don’t

**Dualism**

1. The spirit or soul is independent of the body

A. agree

B. I don’t know

C. disagree

2. The mind or psyche is independent of the brain

A. agree

B. I don’t know

C. disagree

3. María and Juan suffer a car accident and they are taken to the nearest hospital. After an evaluation by the medical staff it is determined that María presents no bodily lesions but is brain dead, while Juan’s brain is unharmed, but he presents bodily lesions incompatible with life. After discussing it with the families and receiving INCUCAI’s [Argentina’s transplant coordination institution] authorization, the neurosurgery team performs an experimental transplant: Juan’s brain in María’s body. Days later the transplanted patient wakes up in the ICU. Whose family would the patient recognize as their own?

A. María’s

B. Juan’s

C. both

D. none of them
