## Additional File 3 for "Knowledge of theory of evolution and beliefs as determinants of dualism in health science students"

Additional File 4.

Table 1. *Results of the Mann Whitney Wilcoxon Tests for the primary outcomes scores between majors.*

**W	p-value**

KEE	10896	0.3438

KEEH	10955	0.2861

Dualism	10182	0.9320

Supernatural beliefs	10560	0.6472

Table 2.

Chi square results for all KEE and KEEH questions by major

| **Question** | **X²** | **p-value** |
| --- | --- | --- |
| KEE 1 | 3.149 | 0.533 |
| KEE 2 | 8.773 | 0.067 |
| KEE 3 | 6.565 | 0.161 |
| KEE 4 | 9.499 | 0.050 * |
| KEE 5 | 1.801 | 0.772 |
| KEE 6 | 13.582 | 0.009 * |
| KEE 7 | 8.644 | 0.071 |
| KEE 8 | 4.061 | 0.398 |
| KEE 9 | 2.697 | 0.747 |
| KEE 10 | 3.133 | 0.536 |
| KEE 11 | 8.007 | 0.091 |
| KEEH 1 | 2.541 | 0.637 |
| KEEH 2 | 2.057 | 0.725 |
| KEEH 3 | 8.571 | 0.073 |

Chi square results for all dualism questions by major

| **Question** | **X²** | **p-value** |
| --- | --- | --- |
| Spirit / body | 1.311 | 0.519 |
| Mind / brain | 0.565 | 0.754 |
| Car accident case scenario | 5.194 | 0.074 |

Chi square results for all religion questions by major

| **Question** | **X²** | **p-value** |
| --- | --- | --- |
| Belief in God | 0.370 | 0.831 |
| Belief in the soul | 1.695 | 0.428 |
| Belief in energies | 2.935 | 0.230 |
| Belief in afterlife | 1.512 | 0.470 |
| Relationship to God | 0.852 | 0.653 |
| Religion | 0.497 | 0.780 |
