## Additional File 4 for "Knowledge of theory of evolution and beliefs as determinants of dualism in health science students"

Additional File 5. *Results for the Principal Components. Proportion of explained variance and loadings for all variables are included.*

**% Variance	KEE	KEEH	Supernatural beliefs	Dualism**

PC1	38.38	0.50	0.48	-0.53	-0.48

PC2	22.33	0.43	-0.55	-0.32	-0.64

PC3	20.00	-0.74	0.67	-0.03	-0.06

PC4	19.29	0.13	0.13	0.78	-0.59
