## Additional File 5 for "Knowledge of theory of evolution and beliefs as determinants of dualism in health science students"

Additional File 6. Results of logistic and multinomial regressions. Signif. codes: 0 ‘***’ 0.001 ‘**’ 0.01 ‘*’ 0.05 ‘.’ 0.1 ‘ ’ 1

```
KEE1
Coefficients:
                      Estimate Std. Error z value Pr(>|z|)
(Intercept)             1.1212     0.4003   2.801  0.00509 **
GenderMale         	0.1605     0.4151   0.387  0.69911
GenderOther             14.0022   974.5970   0.014  0.98854
MajorPsy      		-0.4914     0.3763  -1.306  0.19167
ReligionOther   		0.3077     0.6430   0.479  0.63227
NoReligion    		1.4667     0.3758   3.903  9.5e-05 ***
---

    Null deviance: 231.79  on 286  degrees of freedom
Residual deviance: 212.60  on 281  degrees of freedom
AIC: 224.6


KEE2
Coefficients:
                      Estimate Std. Error z value Pr(>|z|)
(Intercept)             0.4750     0.3187   1.491   0.1360
GenderMale        		-0.5414     0.2681  -2.020   0.0434 *
GenderOther            -16.6243   938.8884  -0.018   0.9859
MajorPsy       		0.2189     0.2497   0.877   0.3806
ReligionOther   		1.0655     0.6931   1.537   0.1242
NoReligion   		-0.3670     0.3001  -1.223   0.2213
---

    Null deviance: 395.32  on 286  degrees of freedom
Residual deviance: 378.56  on 281  degrees of freedom
AIC: 390.56

KEE3
Coefficients:
                      Estimate Std. Error z value Pr(>|z|)
(Intercept)            -0.4662     0.3175  -1.468  0.14204
GenderMale         	0.2921     0.2778   1.052  0.29291
GenderOther            -14.6257   613.0958  -0.024  0.98097
MajorPsy       		0.3731     0.2544   1.467  0.14251
ReligionOther 		 -0.1431     0.5595  -0.256  0.79815
NoReligion    		0.8104     0.2975   2.724  0.00645 **
---

    Null deviance: 388.02  on 286  degrees of freedom
Residual deviance: 371.94  on 281  degrees of freedom
AIC: 383.94

KEE4
Coefficients:
                      Estimate Std. Error z value Pr(>|z|)
(Intercept)            0.09066    0.33089   0.274   0.7841
GenderMale        		0.38156    0.30968   1.232   0.2179
GenderOther            -0.85453    1.44516  -0.591   0.5543
MajorPsy      		0.66730    0.27913   2.391   0.0168 *
ReligionOther  		0.23658    0.58307   0.406   0.6849
NoReligion   		0.62385    0.31703   1.968   0.0491 *

    Null deviance: 335.80  on 286  degrees of freedom
Residual deviance: 325.34  on 281  degrees of freedom
AIC: 337.34

KEE5
Coefficients:
                        Estimate Std. Error z value Pr(>|z|)
(Intercept)              1.23288    0.37494   3.288  0.00101 **
GenderMale         	-0.28031    0.33060  -0.848  0.39651
GenderOther              14.12273 1022.15714   0.014  0.98898
MajorPsy       		-0.04495    0.31323  -0.144  0.88589
ReligionOther    		0.02192    0.64685   0.034  0.97296
NoReligion     		0.47918    0.35131   1.364  0.17258
---

    Null deviance: 277.55  on 286  degrees of freedom
Residual deviance: 274.02  on 281  degrees of freedom
AIC: 286.02

KEE6
Coefficients:
                      Estimate Std. Error z value Pr(>|z|)
(Intercept)             0.5588     0.3234   1.728   0.0839 .
GenderMale         	0.1890     0.2848   0.664   0.5069
GenderOther            -15.9193   941.2385  -0.017   0.9865
MajorPsy      		-0.6543     0.2611  -2.506   0.0122 *
ReligionOther 		-0.3666     0.5598  -0.655   0.5125
NoReligion    		0.4614     0.3018   1.529   0.1263

    Null deviance: 378.04  on 286  degrees of freedom
Residual deviance: 361.44  on 281  degrees of freedom
AIC: 373.44

KEE7
Coefficients:
                       Estimate Std. Error z value Pr(>|z|)
(Intercept)            -0.20841    0.31354  -0.665    0.506
GenderMale        		-0.06919    0.26913  -0.257    0.797
GenderOther             15.05273  620.81672   0.024    0.981
MajorPsy      		-0.11967    0.24838  -0.482    0.630
ReligionOther  		-0.32270    0.57117  -0.565    0.572
NoReligion   		-0.10800    0.29561  -0.365    0.715

    Null deviance: 388.02  on 286  degrees of freedom
Residual deviance: 383.84  on 281  degrees of freedom
AIC: 395.84

KEE8
Coefficients:
                      Estimate Std. Error z value Pr(>|z|)
(Intercept)            0.20389    0.31725   0.643  0.52043
GenderMale       	   -0.08791    0.28351  -0.310  0.75652
GenderOther            0.31097    1.47722   0.211  0.83327
MajorPsy     	   -0.13837    0.25978  -0.533  0.59429
ReligionOther  	    0.09402    0.54643   0.172  0.86340
NoReligion  		   -0.98537    0.29895  -3.296  0.00098 ***
---

    Null deviance: 374.69  on 286  degrees of freedom
Residual deviance: 360.47  on 281  degrees of freedom
AIC: 372.47

KEE9
Coefficients:
                       Estimate Std. Error z value Pr(>|z|)
(Intercept)            -0.40753    0.31530  -1.293    0.196
GenderMale       	    -0.02558    0.26607  -0.096    0.923
GenderOther           -14.40294  624.02418  -0.023    0.982
MajorPsy      	    -0.03105    0.24582  -0.126    0.899
ReligionOther           0.31069    0.55324   0.562    0.574
NoReligion              0.20900    0.29735   0.703    0.482

    Null deviance: 392.55  on 286  degrees of freedom
Residual deviance: 389.66  on 281  degrees of freedom
AIC: 401.66

KEE10
Coefficients:
                      Estimate Std. Error z value Pr(>|z|)
(Intercept)            -0.8734     0.3363  -2.597  0.00941 **
GenderMale        	    -0.0468     0.2734  -0.171  0.86409
GenderOther           -14.3214   623.2324  -0.023  0.98167
MajorPsy      	    -0.1122     0.2522  -0.445  0.65652
ReligionOther   	     0.8232     0.5649   1.457  0.14506
NoReligion             0.5459     0.3192   1.710  0.08719 .
---
    Null deviance: 380.12  on 286  degrees of freedom
Residual deviance: 374.28  on 281  degrees of freedom
AIC: 386.28

KEE11
			Score=1 Estimate	p-val		Score=2 Estimate	p-val
(Intercept)		0.98			0.1047		2.07			0.0001
GenderMale		0.41			0.4636		0.02			0.9611
GenderOther		-3.08			0.7238		10.33			0.9700
MajorPsy		-0.42			0.4307		-0.95			0.0473*
ReligionOther	-1.88			0.1276		-0.04			0.9622
NoReligion		0.26			0.6194		0.87			0.0710


KEEH1
Coefficients:
                        Estimate Std. Error z value Pr(>|z|)
(Intercept)             0.259616   0.314220   0.826    0.409
GenderMale        	     -0.004663   0.271222  -0.017    0.986
GenderOther            13.878396 621.450469   0.022    0.982
MajorPsy      	     -0.048251   0.251032  -0.192    0.848
ReligionOther   	     0.642337   0.592546   1.084    0.278
NoReligion    	     0.270474   0.295272   0.916    0.360


    Null deviance: 382.08  on 286  degrees of freedom
Residual deviance: 378.59  on 281  degrees of freedom
AIC: 390.59

KEEH2
Coefficients:
                      Estimate Std. Error z value Pr(>|z|)
(Intercept)           -1.14780    0.34689  -3.309 0.000937 ***
GenderMale            0.45649    0.28057   1.627 0.103733
GenderOther           1.34521    1.48595   0.905 0.365312
MajorPsy              0.09349    0.26627   0.351 0.725516
ReligionOther 	   -0.70365    0.71789  -0.980 0.327008
NoReligion   	    0.21533    0.32335   0.666 0.505460
---

    Null deviance: 353.80  on 286  degrees of freedom
Residual deviance: 348.05  on 281  degrees of freedom
AIC: 360.05

KEEH3
Coefficients:
                      Estimate Std. Error z value Pr(>|z|)
(Intercept)             0.6410     0.3250   1.972   0.0486 *
GenderMale              0.1938     0.2854   0.679   0.4970
GenderOther            -0.4765     1.4749  -0.323   0.7466
MajorPsy               -0.4488     0.2596  -1.728   0.0839 .
ReligionOther          -0.1703     0.5541  -0.307   0.7586
NoReligion             0.2900     0.3038   0.955   0.3398
---

    Null deviance: 369.81  on 286  degrees of freedom
Residual deviance: 363.64  on 281  degrees of freedom
AIC: 375.64


Dualism1
			Disagree Estimate	p-val		Don’t know Estimate	p-val
(Intercept)		-0.13			0.7255		-0.61			0.0001
GenderMale		0.28			0.4341		0.53			0.1565
GenderOther		-11.23			0.9546		0.12			0.9341
MajorPsy		0.39			0.2173		0.20			0.5465
ReligionOther	0.40			0.5232		-0.29			0.7130
NoReligion		0.79			0.0251*	1.06			0.0072**


Dualism2
			Disagree Estimate	p-val		Don’t know Estimate	p-val
(Intercept)		2.06			>0.001		-0.19			0.8068
GenderMale		-0.75			0.0538		-2.97			0.0067**
GenderOther		9.03			0.9618		-3.76			0.5253
MajorPsy		0.08			0.8301		-0.34			0.5604
ReligionOther	-0.05			0.9492		0.73			0.5534
NoReligion		0.19			0.6587		0.60			0.4321


Dualism3
			María’s Estimate	p-val		Don’t know Estimate	p-val
(Intercept)		0.52			0.0042		0.38			0.0013
GenderMale		0.77			0.0590		0.32			0.3417
GenderOther		37.55			0.8967		1.46			0.2690
MajorPsy		0.48			0.2172		0.31			0.0760
ReligionOther	1.11			0.2339		0.74			0.2565
NoReligion		0.48			0.0313*	0.35			0.1935
```
